# A Common Neural Signal of Evidence Accumulation for Perceptual and Mnemonic Decisions

**DOI:** 10.1101/2025.11.13.688140

**Authors:** Alice Tsvinev, April Pilipenko, Hannah Hausman, Jason Samaha

## Abstract

Humans frequently make decisions based on sensory input from the external environment or information retrieved from memory. The centro-parietal positivity (CPP), an event-related EEG potential, has recently been identified as a neural correlate of sensory evidence accumulation during perceptual decision-making tasks. However, it remains unclear whether this component also reflects the accumulation of evidence in service of decisions grounded in semantic and episodic long-term memory. Across two experiments, we investigated whether the CPP serves as a domain-general neural signal of evidence accumulation. In Experiment 1, participants completed 2AFC perceptual and semantic memory tasks with varying levels of evidence strength. Perceptual judgements involved luminance discrimination of alphanumeric strings with three luminance difference levels controlling perceptual evidence strength. Semantic memory judgements involved discriminating population differences between U.S. states with census data used to define three bins of memory evidence strength. A CPP component was observed in both tasks whose build-up rate (i.e., slope) scaled with evidence strength, response time, and confidence in both stimulus-and response-locked analyses. Extending these findings to episodic memory, participants in Experiment 2 completed a two-alternative forced-choice word recognition task with target words varying in exposure frequency during learning to control episodic memory strength. Again, we found that CPP slopes scaled with memory strength, response time, and confidence. Together, these findings support the CPP as a domain-general neural signature of evidence accumulation across perceptual, semantic, and episodic mnemonic decisions.

## Introduction

Humans frequently make decisions based on sampling sensory evidence from our external environment but also by internally sampling and retrieving information held in memory (O’Connell et al., 2012; Shadlen & Shohamy, 2016; Van Ede & Nobre, 2024). In recent decades, cognitive neuroscience has made significant strides in understanding how the brain supports this process, revealing that many decisions are made by accumulating information over time until a decision threshold is reached. Indeed, research spanning computational modelling of human behavior to direct neural recordings from behaving animals supports accumulation-to-bound as a mechanistic explanation for a range of decision behaviors including variation in response times (RT), accuracy, and confidence (Desender et al., 2021; Kiani et al., 2014; O’Connell et al., 2018; Ratcliff et al., 2016; Ratcliff & McKoon, 2008).

Recent work in humans has uncovered an electroencephalogram (EEG) potential that closely tracks the sensory evidence accumulation process underlying simple perceptual decisions (O’Connell et al., 2012). The centro-parietal positivity (CPP) event-related potential (ERP) has been found to closely follow the expected temporal dynamics of sensory evidence accumulation during perceptual decisions: it gradually build ups in proportion to the strength of the unsigned sensory evidence, peaks around the time of behavioral response, and can predict decision accuracy, RT, and confidence in simple perceptual decisions (Charles et al., 2020; Dou et al., 2024; Herding et al., 2019; Kelly & O’Connell, 2013; O’Connell et al., 2012). Furthermore, it has been argued that the CPP reflects a supramodal decision signal, occurring with a similar topography and temporal dynamic regardless of the sensory modality (e.g. auditory or visual) or response format (e.g., button press or covert response such as mental counting), indicative of a domain-general decision-making process in the human brain (O’Connell et al., 2012; Schaffhauser et al., 2021).

Despite the substantial literature linking the CPP to evidence accumulation, most studies have examined the CPP selectively in the context of perceptual decision-making. Although several recent studies have observed a CPP-like component during working-memory based decisions (Van Ede & Nobre, 2024; van Vugt et al., 2019), whether the CPP tracks decisions made on the basis of accumulating evidence from long term memory remains unknown. This raises an important hypothesis regarding the claim of domain-generality of the CPP: if the underlying neural dynamics reflect a central decision-making mechanism, the CPP should track evidence accumulation for both sensory and mnemonic evidence and be predictive of decision behaviors in both cases.

Across two experiments, we tested this hypothesis by directly comparing perceptual and mnemonic decision dynamics across both semantic and episodic memory tasks with varying levels of evidence strength. In a first experiment, we compared the neural dynamics of decision-making in a perceptual luminance discrimination task to a discrimination task involving retrieval of semantic knowledge about US state populations. In a second experiment, we employed a two-alternative forced-choice (2AFC) word recognition task to examine the neural dynamics underlying decisions based in episodic memory. We examined whether a CPP-like component emerges across all tasks, scales similarly with evidence strength, and predicts decision behaviors like RT and confidence. We predicted that a CPP with similar topography would emerge across tasks and that the slope of the CPP would increase with stronger sensory and mnemonic evidence, faster RTs, and higher subjective confidence, regardless of the task.

## Methods

### Participants

24 participants (15 female, 8 male, 1 nonbinary; *M*_age_ = 21 years, *SEM*_age_ = 0.18 years) completed Experiment 1, and 30 participants (18 female, 8 male, 1 nonbinary, 1 genderfluid, 2 undisclosed; *M*_age_ = 21 years, *SEM*_age_ = 0.62 years) completed Experiment 2. All participants were recruited from the University of California, Santa Cruz (UCSC) in exchange for course credit, reported normal or corrected-to-normal vision, and provided written informed consent. All procedures performed in this study were reviewed and approved by the institutional review board at the University of California, Santa Cruz.

We applied a consistent exclusion criteria across the two experiments leading to the exclusion of one participant due to corrupted EEG data and three participants due to low accuracy (below 60%, where chance is 50%) in the easiest difficulty condition of the mnemonic task in Experiment 1, suggesting little familiarity with US state population sizes. One participant was removed from analysis due to excessive noise (e.g., >35% trials rejected) and three were removed due to low accuracy (below 60%) in the highest exposure (i.e., easiest) condition in Experiment 2. Thus, a final sample of 20 participants and 26 participants were involved in analysis for Experiment 1 and Experiment 2, respectively. The sample size for Experiment 1 was chosen to be on par with our recent work on perceptual evidence accumulation (Dou et al., 2024; Morrow et al., 2024). A power analysis of the results from Experiment 3 of Dou et al. (2024), who reported effect sizes for the effect of evidence strength on stimulus-and response-locked CPP slopes, suggests that 5 to 18 subjects, respectively, are needed to detect an effect with 80% power. A power analysis of the results on the effects of evidence strength on stimulus-and response-locked CPP slopes in the semantic memory task from Experiment 1, suggests that 19 to 24 subjects are needed, respectively, to detect an effect with 80% power, resulting in the selected sample size for Experiment 2.

### Stimuli and Apparatus

In all experiments, stimuli were presented on a grey background (∼30 cd/m^2^) using an electrically shielded VIEWPixx/EEG monitor (120 Hz refresh rate, resolution 1920 × 1080 pixels) that was ∼53 cm wide and was viewed at a distance of either ∼70 cm (Experiment 1) or ∼75 cm from a chinrest (Experiment 2). Stimulus presentation and behavioral data were controlled by PsychToolbox (Version 3; Kleiner et al., 2007; Pelli, 1997) running in the MATLAB environment.

#### Experiment 1

The perceptual task consisted of two strings of eight capitalized ‘X’ characters (Helvetica font, size 16) presented directly (0.3 degrees of visual angle) above and below the central fixation point. One string was designated the “comparison” stimulus as it varied in luminance intensity across three levels (34.3, 34.8, 37 cd/m^2^), randomly determined on each trial. The luminance of the other “standard” stimulus was fixed at 33.9 cd/m^2^. The difference in luminance between the standard and comparison string determined the strength of evidence for the perceptual decision. In the semantic memory task, the two strings were the names of states in the U.S. and were presented at the same locations above and below the central fixation point, with each string having the same luminance value (33.9) for all trials. The strength of memory evidence was manipulated by creating state population pairs that varied in their degree of population differences. To this end, we took 2020-2024 US census data (US Census Bureau, 2024) for each US state and calculated the absolute value of population differences for every possible state combination. Three difficulty levels were defined based on the absolute differences between state pairs, chosen based on behavioral pilot data: Hard (20-45th percentile), Medium (45-80th percentile), and Easy (80-100th percentile). This partitioning let 306 state pairs fall into the Hard bin, 429 state pairs fall into the Medium bin, and 245 state pairs fall into the Easy bin. Each trial randomly sampled (with replacement) a state pair from all pairs falling within the assigned difficulty bin, ensuring a balanced number of trials per difficulty level. This manipulation provided a quantifiable form of evidence strength comparable to the difficulty manipulation in luminance values in the perceptual task.

#### Experiment 2

Stimulus presentation parameters matched those of Experiment 1 unless otherwise noted. The 2AFC word recognition task consisted of separate learning and test phases. During learning, centrally presented word strings were shown 1, 2, or 4 times to manipulate episodic memory strength. Following a 60-second retention interval, pairs of learned words and non-learned foils were presented simultaneously above and below fixation. All stimuli were presented at the same luminance used in the semantic memory task of Experiment 1 (33.9 cd/m^2^). A master list of 672 word stimuli were selected from the SUBTLEXus database (Brysbaert et al., 2014) to control for word frequency (rating of 1.68–2.00; 70–80th percentile) concreteness (rating of 2.5–3.44; 45–65th percentile), and word length (5-7 letters per word). Subject-specific lists of target (i.e., to-be-learned) words and foils were generated for each participant by random selection from the master list and subsequently divided into three blocks by pseudorandom selection without replacement, with to-be-learned items further divided into the three exposure frequency conditions.

To minimize ERP responses caused by transient stimulus onsets, stimuli linearly ramped up in luminance across the first 300 ms (Experiment 1) or 350 ms (Experiment 2) of presentation time and remained on the screen until a response was given or the trial timed out (see Procedure).

### Procedure

#### Experiment 1

Participants were tested individually in a dim, sound-attenuated room. The experimenter explained the instructions to the participant and verified that the participant understood the instructions before proceeding to the practice trials and critical blocks. The participant completed four practice blocks (20 trials each) and eight critical blocks (80 trials each) across the perceptual and semantic memory task with task order counterbalanced across participants. Across all trials, participants heard a beep if they did not respond within the designated response window (1.8 seconds for perceptual; 3 seconds for semantic memory), prompting them to continue to a confidence response without a decision response.

In the perceptual task, participants were informed that two strings of X’s would gradually fade into the screen above and below the central fixation point, and their task was to select the brighter string. For the semantic memory task, participants were asked to identify which of the two presented US states had a greater population. No training regarding state population information was given prior to beginning the task. In each task, participants were instructed to respond as quickly and accurately as possible and then rate their confidence on a four-point scale. Stimuli remained visible until a response was made (or until the response window timed out), after which confidence ratings were collected.

For half the blocks, participants were instructed to keep their right index and middle fingers on the keys “J” and “K” representing the location of the target stimulus being either above or below the central fixation, respectively. Participants were additionally instructed to keep their left-hand fingers on the keys “1”, “2”, “3”, and “4” (from pinky to index finger) to indicate confidence from lowest to highest, respectively. To minimize lateralized motor-related activity in the averaged ERP signal, the response-hand mapping was switched every other block (with the order of response mappings counter balanced across subjects). In blocks where the response mapping was flipped, participants were instructed to keep their left index and middle fingers on the keys “F” and “D” representing the location of the target stimulus being either below or above the central fixation, respectively. Participants were additionally instructed to keep their right-hand fingers on the right keys “7”, “8”, “9”, and “0” (from pinky to index finger) to indicate confidence from lowest to highest, respectively.

Each participant completed 320 trials per task (640 total), with difficulty levels randomized across trials.

#### Experiment 2

Participants completed a 2AFC word recognition task consisting of separate learning and test phases. Participants completed two practice blocks (84 learn; 12 test trials each) and four critical blocks (196 learn; 84 test trials each).

During learning, centrally presented words were shown serially for 1150ms with jittered inter-stimulus intervals (400-700ms). Words were presented according to their assigned exposure condition (4x, 2x, 1x) in four subphases that participants were unaware of. To manipulate mnemonic strength while minimizing serial-position effects, words from the 4x exposure condition were presented across all four subphases, words from the 2x exposure condition were presented in only the last two subphases, and words from the 1x exposure group were only presented in the last subphase of learning.Participants were instructed to pay attention to and learn the words for later testing, and no responses were required during learning. Following a 60-second retention interval, participants completed the test phase, in which pairs consisting of one learned word and one non-learned foil were presented above and below fixation. Participants were tasked with identifying the previously learned word within a 2.3 second response window, followed by an untimed confidence rating. Each participant completed 784 learning trials and 336 testing trials total. All other procedures, such as hand response mappings and trial time-out beeps, were identical to Experiment 1.

### EEG Recording and Analysis

In all experiments, continuous EEG was acquired from 63 active electrodes (BrainVision actiCHamp), with impedance at each central-parietal electrode kept below 20kΩ. Recordings were digitized at 1000 Hz, and FCz was used as the online reference. EEG was processed offline using custom scripts in MATLAB (version R2019b) and the EEGLAB toolbox (Delorme & Makeig, 2004). Data were high-pass filtered at 0.1 Hz and low-pass filtered at 30 Hz, then downsampled to 500 Hz. Data were re-referenced offline to the average of all electrodes. Continuous signals were then segmented into epochs centered on stimulus onset using distinct time windows for the perceptual (-1,000ms to 2,500ms; Experiment 1), semantic memory (-1,000ms to 3,200ms; Experiment 1), and episodic memory (-1,000ms to 2,500ms; Experiment 2) tasks. Individual trials were rejected if any scalp channel exceeded ±100 μV at any time during the interval extending from-300ms to 1000ms relative to stimulus onset. On average, 36 trials were rejected per subject across both tasks of Experiment 1, and 15 trials were rejected per subject in Experiment 2. In all experiments, trials excluded from the EEG data were similarly excluded from the analyses of behavior. Noisy channels were interpolated, and an independent components analysis was performed to identify and remove components reflecting eye blinks or movements. On average, 1.2 components were removed per subject in Experiment 1, and 1 component was removed per subject in Experiment 2 (range = 1-2). Lastly, a prestimulus baseline of-200ms to 0ms was subtracted from each trial in both experiments.

## Statistical Analyses

In both experiments, the CPP buildup rate was defined as the slope of a line fit to each participants’ average CPP waveform. The CPP slope was the main focus of our EEG analysis as it is most theoretically related to the construct of evidence-accumulation rate, with steeper slopes reflecting a faster build-up of evidence (Kelly & O’Connell, 2013; O’Connell et al., 2012). For each condition, EEG data were averaged across trials and over a predefined set of centro-parietal electrodes (CP1, CP2, CP3, CP4, CPz, P1, P2, P3, P4, Pz), yielding a single ERP waveform per subject per condition.

In both experiments, we computed the slope of the CPP component by fitting a line to the ERP waveform for each subject, task (perceptual, semantic memory; episodic memory) and difficulty/exposure condition (easy, medium, hard; 4x, 2x, 1x). For the response-locked analyses, the slopes were calculated over time windows between-800ms to-100ms in the perceptual task,-1200ms to-100ms in the semantic task, and-600 to-100ms in the episodic task, all relative to the response. For the stimulus-locked analyses, the slopes were calculated over time windows between 300ms to 700ms (perceptual), 300ms to 1000ms (semantic), and 650ms to 1000ms (episodic). For all experiments, these time windows were determined based on visual inspection of the grand-average ERP from each task separately (blind to the difficulty level) to best capture the rising phase of the stimulus-and response-locked CPP waveform. For Experiment 2, only EEG data during testing was analyzed.

In Experiment 1, we conducted exploratory correlational analyses (as our sample size was not chosen to examine individual differences) to test if participants might use a common threshold for terminating perceptual and mnemonic decisions. If the CPP reflects domain-general bounded evidence accumulation, then the mean amplitude of the CPP just prior to the response should reflect the amount of evidence a given participant requires before committing to a choice. To test for a common threshold across tasks, we computed the Spearman correlation across participants between the mean pre-response (-100 to-25ms) CPP amplitudes in the memory and perceptual tasks. We next considered response times, which within evidence accumulation frameworks are jointly influenced by the drift rate and decision boundary. While we would not necessarily expect participants who accumulated perceptual evidence quickly during luminance judgements to also accumulate semantic memory evidence quickly during state population judgements, a common decision boundary may generalize across tasks. If participants employ a common decision boundary across tasks, this shared decision policy may produce correlated response times despite task-specific differences in drift rate. We therefore also computed Spearman correlations between median RTs in each task (averaged across difficulty levels) to assess whether response times were consistent with a shared decision boundary across tasks.

Finally, we conducted additional analyses in all experiments to investigate if the CPP slope varied as a function of both response time and confidence within each difficulty/exposure level. For each subject, task, and difficulty level, trials were split into fast and slow RTs based on a median split, and into high and low confidence trials based on a mean split, consistent with prior work linking CPP dynamics to decision timing and subjective certainty (Dou et al., 2024; Herding et al., 2019; Kurtz et al., 2017). A line was fit to the resulting CPP waveforms using the same time windows as the previous analysis.

## Results

### A Neural Signature of Evidence Accumulation Across Perception and Semantic Memory

We sought to identify whether the CPP extends beyond perceptual decisions and indexes a domain-general neural decision variable. To this end, we manipulated the quality of perceptual and semantic memory evidence and tested the CPP for three features consistently observed in prior work (Kelly & O’Connell, 2013; O’Connell et al., 2012). Namely, if the CPP reflects a domain-general neural index of evidence accumulation, its buildup rate (quantified by slope) should 1) increase as the quality of evidence increases, 2) increase as RTs decrease, and 3) increase as subjective confidence increases, regardless of evidence source. In Experiment 1, perceptual and semantic evidence strength were manipulated across three difficulty levels using luminance differences and US state population differences, respectively (Figure 1A).

**Figure 1.**
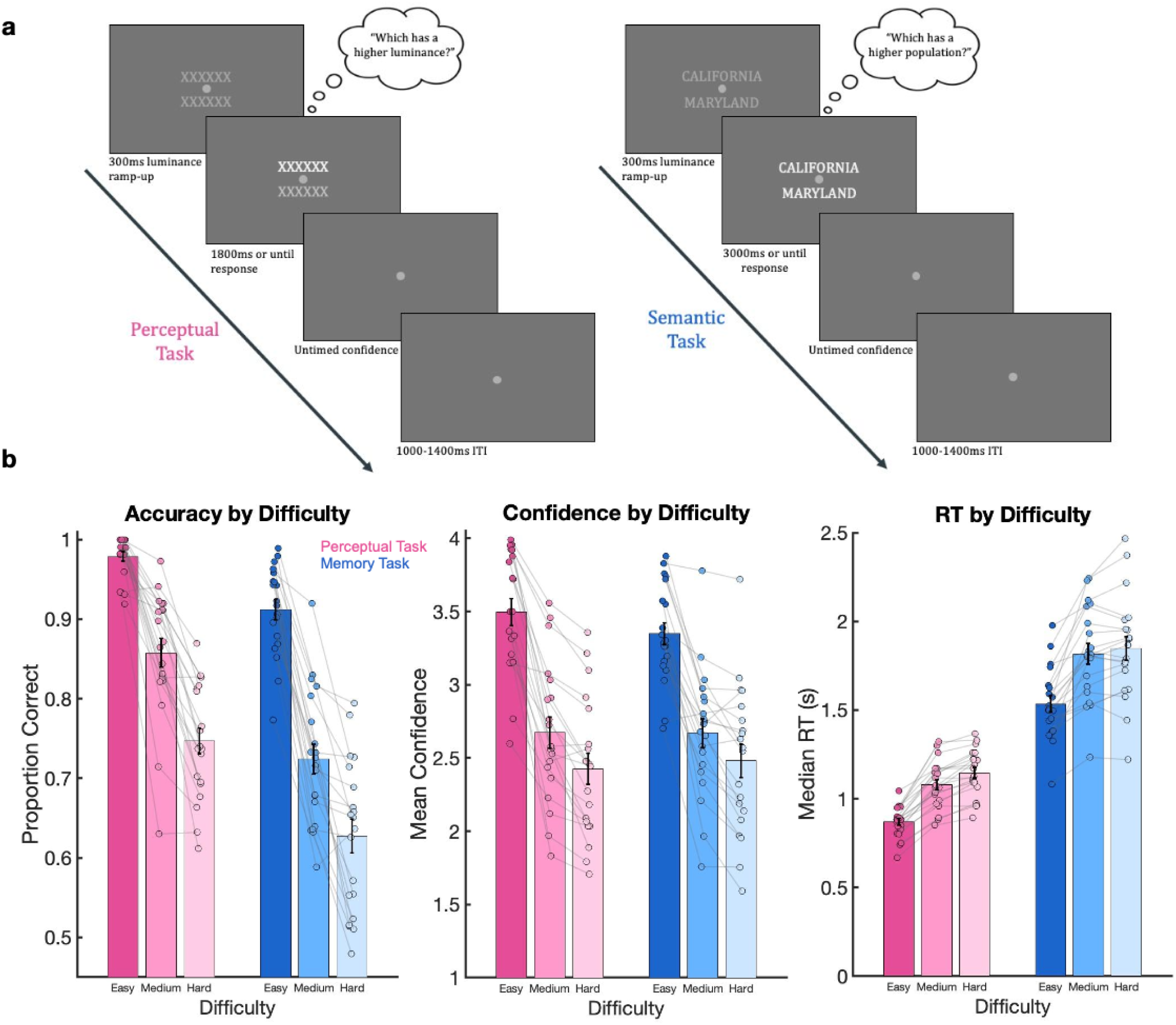
Example trials and behavioral results for Experiment 1 (N = 20). A) Each trial began with a central fixation dot and a 300ms gradual luminance ramp up of the perceptual (top left) or semantic memory (top right) stimuli, which were displayed until response (up to a maximum of 1800 or 3000 ms, respectively). Stimuli were presented at one of three difficulty levels randomly for each trial. Participants were tasked with choosing the brighter alphanumeric string (perceptual) or more populated state (semantic memory) and rating their confidence in that decision (on a scale of 1-4). B) As difficulty increased in both tasks, mean accuracy (left plot) and confidence ratings (middle plot) decreased, and median reaction time (right plot) increased. Error bars denote ±1 SEM (across participants). Colored dots over each bar represent individual subject scores per measure, with gray lines connecting scores across all difficulty levels per subject.

At the behavioral level, three 2 (task: perceptual, memory) x 3 (difficulty: easy, medium, hard) repeated-measures ANOVAs were conducted to examine the effects of task type and difficulty on accuracy, reaction time, and subjective confidence. As expected, increasing difficulty led to lower accuracy, *F*(2, 38) = 419.66, *p* =1.25×10^-26^; lower confidence, *F*(2, 38) = 99.41, *p* = 7.97×10^-16^; and slower RTs, *F*(2, 38) = 84.94, *p* = 9.50×10^-15^, across both tasks (Figure 1B).

Performance additionally differed across tasks, with perceptual decisions yielding greater accuracy, *F*(1, 19) = 23.77, *p* = 1.05×10^-04^; and faster RTs, *F*(1, 19) = 250.75, *p* = 2.11×10^-12^; relative to semantic decisions. However, no significant difference in overall confidence between perceptual and semantic decisions was observed, *F*(1,19) = 0.144, *p* =.708, despite notable differences in performance and RT. Interestingly, an interaction between task and difficulty was observed on confidence, *F*(2,38) = 3.52, *p* =.040, with greater confidence ratings for the perceptual task than the semantic memory task, but only among the easy trials. No other interactions were observed between task and difficulty on either accuracy, *F*(2, 38) = 2.78, *p* =.075; or RT, *F*(2,38) = 2.02, *p* =.146.

During decision formation, we observed a positive ERP component over central parietal electrodes in the perceptual task which peaked around 650 ms poststimulus (Figure 2A, left) and ramped up prior to motor response (Figure 2A, right), resembling the CPP (O’Connell et al., 2012). A similar positive ERP component was observed in the semantic memory task, peaking around 1000 ms poststimulus (Figure 2B, left) and ramping up prior to the motor response (Figure 2B, right). We then correlated the CPP amplitudes at each electrode between the perceptual and semantic memory tasks to quantify the similarity of their scalp topographies. Notably, the resulting topographies were highly similar for both stimulus-(r = 0.93, *p* <.001) and response-locked (r = 0.959, *p* <.001) signals, indicating that the CPP exhibited a similar spatial distribution across tasks. Although the CPP peaked later in the semantic memory task, consistent with longer response times observed in that task, its scalp distribution remained highly similar. Together, these findings suggest that decisions made using perceptual and semantic memory information may engage a common neural process, providing initial evidence for a domain-general neural signature of evidence accumulation.

**Figure 2.**
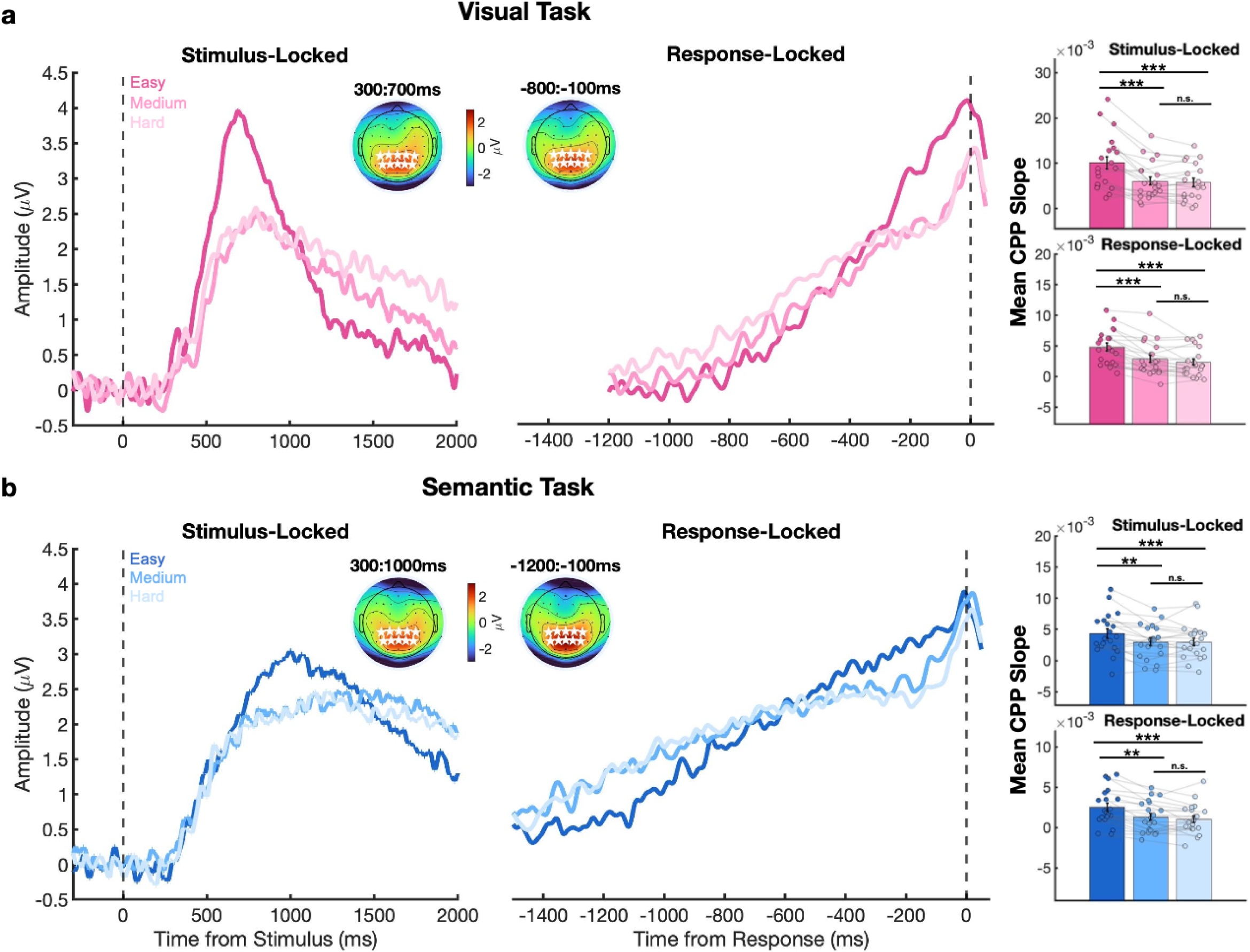
Effects of perceptual and semantic memory task difficulty on the CPP component in Experiment 1 (N = 20). Grand average CPP waveforms were aligned to both stimulus onset (left column) and response execution (right column). The vertical dashed line denotes the time of stimulus onset (left) and time of response execution (right). The inset topographies of the ERP measured after stimulus onset for the visual perception task (A) and for the semantic memory task (B) show a positive central-parietal component (electrodes used for analysis are shown in white stars) across both stimulus and response-locked windows. Bar plots (far right) show mean CPP slopes per difficulty in each task, with individual subjects plotted as points and the connecting light gray lines depicting individual slope trends across difficulty. Note that the topography time windows are different to account for the longer memory-based decision time. Stimulus-aligned and response-aligned CPP slopes both increased significantly as evidence strength increased for both the perception and memory tasks.

To test if the CPP was sensitive to the strength of memory and/or perceptual evidence (and task), we ran two 2 (task: perceptual, semantic memory) x 3 (difficulty: easy, medium, hard) repeated-measures ANOVAs predicting CPP slope separately for response-and stimulus locked waveforms. As shown in Figure 2, CPP slopes were steeper for perceptual relative to semantic decisions in both stimulus-locked (*F*(1,19) = 27.97, *p* = 4.20×10^-05^) and response-locked (*F*(1,19) = 25.17, *p* = 7.65×10^-05^) analyses. Critically, CPP slopes varied as a function of evidence strength, with stronger evidence (easier trials) resulting in steeper buildup rates in both stimulus-locked (*F*(2,38) = 34.57, *p* = 2.79×10^-09^) and response-locked (*F*(2,38) =37.30, *p* = 1.09×10^-09^) analyses. Significant Task x Difficulty interactions emerged for both stimulus-locked (*F*(2,38) =14.38, *p* = 2.24×10^-05^) and response-locked (*F*(2,38) = 3.40, *p* = 0.044) CPP slopes, indicating that while the CPP slope increased with evidence strength in both tasks, the effect of difficulty was more pronounced in the perceptual task than in the semantic memory task. This difference was expected, as no attempt was made to equate the difficulty manipulation across tasks.

Because the Task x Difficulty interaction could be driven by a lack of difficulty effect in one task, we conducted additional one-way repeated measures ANOVAs and repeated-measures *t*-tests predicting CPP slopes from difficulty level within each task separately. If the CPP reflects a domain-general evidence accumulation signal, difficulty effects should be observed on the CPP slope in both tasks. Consistent with this prediction, CPP slopes varied significantly with difficulty in the perceptual task, for both stimulus-locked (*F*(2,38) = 31.16, *p* = 9.76×10^-09^) and response-locked (*F*(2,38) = 26.43, *p* = 6.41×10^-08^) analyses, with steeper slopes for Easy relative to both Hard (stimulus-locked: *t*(19) = 6.32, *p* = 4.58×10^-06^; response-locked: *t*(19) = 5.90, *p* = 1.12×10^-05^) and Medium conditions (stimulus-locked: *t*(19) = 5.46, *p* = 2.88×10^-05^; response-locked: *t*(19) = 6.21, *p* = 5.75×10^-06^). No differences were observed between Hard and Medium trials (stimulus-locked: *t*(19) = 0.98, *p* = 0.337; response-locked: *t*(19) = 1.70, *p* = 0.105). Critically, a similar pattern was observed in the semantic memory task, with significant effects of Difficulty in both stimulus-locked (*F*(2,38) = 11.57, *p* = 1.19×10^-04^) and response-locked (*F*(2,38) = 17.99, *p* = 3.18×10^-06^) analyses. CPP slopes were again steeper for Easy compared to Hard (stimulus-locked: *t*(19) = 4.55, *p* = 2.19×10^-04^; response-locked: *t*(19) = 4.56, *p* = 2.15×10^-04^) and Medium conditions (stimulus-locked: *t*(19) = 3.71, *p* =.001; response-locked: *t*(19) = 5.67, *p* = 1.80×10^-05^), with no difference between Medium and Hard (stimulus-locked: *t*(19) =-0.03, *p* =.978; response-locked: *t*(19) = 1.23, *p* =.234).

### Reaction Time Analyses

To test whether the CPP also reflected decision-related behaviors across both tasks, we next examined whether CPP slopes varied with RT in both the semantic memory and perceptual tasks. Trials were categorized as fast or slow within each participant, task, and difficulty level using a median split on RTs. As shown in Figure 3, CPP slopes varied as a function of RT, with faster decisions associated with steeper slopes across both perceptual (stimulus-locked: *F*(1,19) = 21.81, *p* = 1.67×10^-04^; response-locked: *F*(1,19) = 31.47, *p* = 2.08×10^-05^) and semantic tasks (stimulus-locked: *F*(1,19) = 15.5, *p* = 8.85×10^-04^; response-locked: *F*(1,19) = 31.11, *p* = 2.23×10^-05^). More difficult decisions additionally resulted in shallower CPP slopes across both perceptual (stimulus-locked: *F*(2,38) = 31.24, *p* = 9.48×10^-09^; response-locked: *F*(2,38) = 26.30, *p* = 6.77×10^-08^) and semantic (stimulus-locked: *F*(2,38) = 11.57, *p* = 1.19×10^-04^; response-locked: *F*(2,38) = 18.06, *p* = 3.07×10^-06^) tasks. A significant interaction between RT and Difficulty was observed in the semantic memory task (stimulus-locked: *F*(2,38) = 3.84, *p* =.030; response-locked: *F*(2,38) = 13.51, *p* = 3.70×10^-05^), whereas this interaction was not reliably present in the perceptual task (stimulus-locked: *F*(2,38) = 1.98, *p* =.152; response-locked: *F*(2,38) = 4.71, *p* =.015). In the semantic memory task, the difference in CPP slope between fast and slow responses varied across difficulty levels, being most pronounced for more difficult trials and attenuated for easier trials, whereas the RT-related differences in CPP slope remained relatively stable across difficulty levels in the perceptual task. These results suggest that while the CPP amplitude varied systematically with both RT and difficulty across tasks, the influence of RT depended on difficulty level primarily in the semantic memory task.

**Figure 3.**
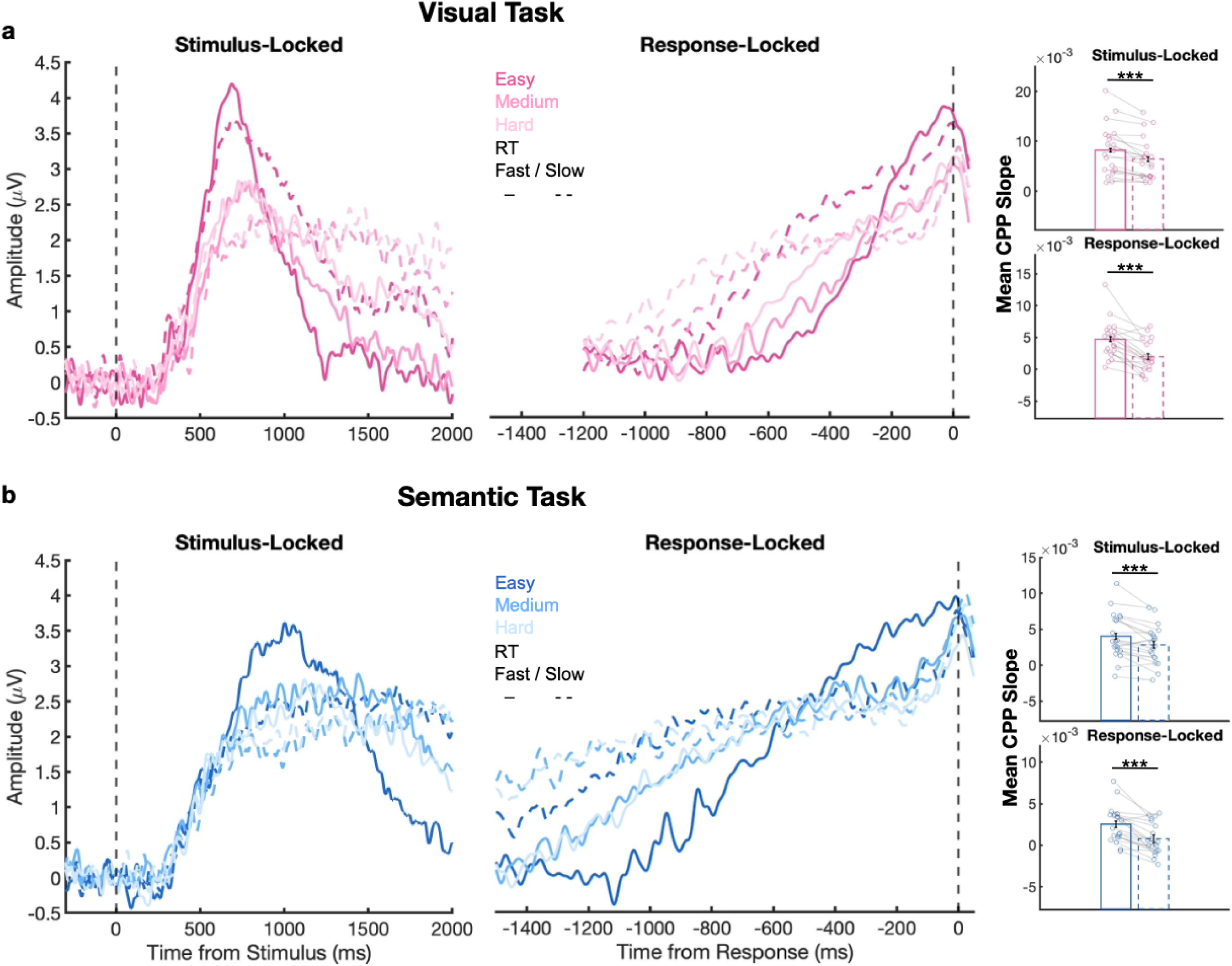
Effects of RT and evidence strength on the CPP in Experiment 1 (N = 20). A) Grand average CPP waveforms aligned to both stimulus onset (left column) and response execution (right column). The vertical dashed line denotes the time of stimulus onset (left) and time of response execution (right). The CPP was broken down by difficulty level for each task and then split into fast (solid line) and slow (dashed line) RT trials within each difficulty level based on a median split. Both stimulus-aligned and response-aligned CPP slopes were steeper on fast-RT trials as compared to slow-RT trials. Bar plots (far right) show mean CPP slopes collapsed across difficulty, split between fast (solid) and slow (dashed) RTs, with individual subjects plotted as points.

### Confidence Analyses

Prior studies have correlated the CPP with subjective confidence judgments, such that higher rates of reported confidence are associated with steeper CPP slopes in perceptual decisions (Dou et al., 2024; Grimaldi et al., 2015; Herding et al., 2019). Following from the hypothesis that the CPP reflects a domain-general signal of evidence accumulation, we predicted that higher subjective confidence would correspond to steeper CPP slopes during both perceptual and semantic memory decisions. Trials were categorized as High or Low Confidence within each participant, task, and difficulty level using a mean split on confidence reports. As shown in Figure 4, CPP slopes were positively correlated with Confidence, such that higher confidence corresponded to steeper CPP slopes across both perceptual (stimulus-locked: *F*(1,19) = 6.03, *p* =.024; response-locked: *F*(1,19) = 12.95, *p* =.002) and semantic memory tasks (stimulus-locked: *F*(1,19) = 5.01, *p* =.037; response-locked: *F*(1,19) = 6.36, *p* =.021). As in previous analyses, CPP slopes also varied as a function of Difficulty across both perceptual (stimulus-locked: *F*(2,38) = 18.64, *p* = 2.29×10^-06^; response-locked: *F*(2,38) = 8.32, *p* =.001) and semantic (SL: *F*(2,38) = 12.19, *p* = 8.13×10^-05^; RL: *F*(2,38) = 14.16, *p* = 2.53×10^-05^) tasks. Notably, an interaction between Confidence and Difficulty was observed in only the response-locked (*F*(2,38) = 9.89, *p* = 3.48×10^-04^) analyses of the semantic memory task, but not in the stimulus-locked (*F*(2,38) = 2.97, *p* =.063) or either perceptual (stimulus-locked: *F*(2,38) = 1.63, *p* =.217; response-locked: *F*(2,38) = 0.488, *p* =.618) analyses. Specifically, in the semantic memory task, the difference in CPP slope between high-and low-confidence trials varied across difficulty levels, being largest for more difficult trials and attenuating for easier trials, whereas the effect of confidence on CPP slope remained relatively stable across difficulty levels in the perceptual task. These results indicate that the influence of confidence on CPP slope depended on task difficulty in the semantic memory decisions, but was comparatively invariant across difficulty levels in perceptual decisions.

**Figure 4.**
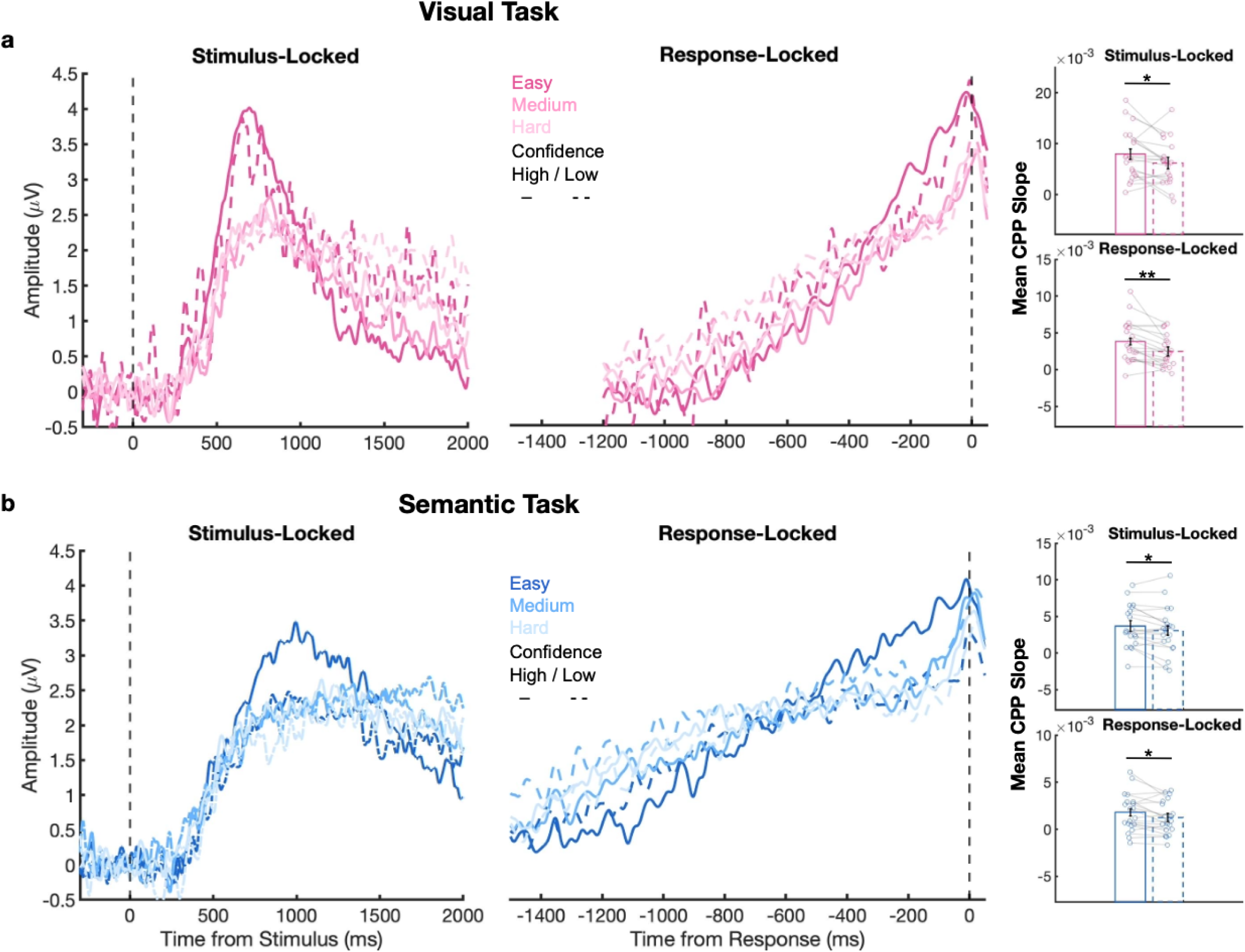
Effects of confidence on the CPP in Experiment 1 (N = 20). A) Grand Average CPP waveforms aligned to both stimulus onset (left column) and response execution (right column). The CPP was broken down by difficulty level for each task and then split into high (solid line) and low (dashed line) Confidence trials within each difficulty level based on a mean split. Both stimulus-aligned and response-aligned CPP slopes were steeper on high-confidence trials compared to low-confidence trials. Bar plots (far right) show mean CPP slopes collapsed across difficulty, split between low (dashed) and high (solid) Confidence, with individual subjects plotted as points.

### Threshold Analyses

In an exploratory analysis examining whether the peak pre-response CPP amplitude reflects a domain-general threshold applied to both perceptual and semantic memory decisions, we correlated each subject’s pre-response (-100 to - 25 ms) response-locked CPP amplitude in the perceptual and semantic memory tasks. We predicted that individuals who exhibit larger pre-response peak CPP amplitude (averaged over difficulty level) in one task would also have a larger amplitude in the other task. Consistent with this prediction, pre-response peak CPP amplitudes were strongly correlated across perceptual and semantic memory tasks, (ρ = 0.78, p = 5.25×10^-5^), indicating that peak pre-response CPP amplitude could reflect a stable signature of the threshold of required evidence which generalizes across domains (Figure 5A). Within the modeling framework, response times reflect both task-specific drift rates and potentially shared decision boundaries, which may lead to correlated response times across tasks. As such, we further predicted that individuals who exhibited faster RTs in one task would exhibit faster RTs in the other task. Consistent with this prediction, a medium positive correlation was observed between median reaction times (averaged over difficulty level) in the perceptual task and semantic memory task (ρ = 0.43, *p* = 0.058), suggesting a trend towards a common underlying factor influencing decision speed across domains (Figure 5B).

**Figure 5.**
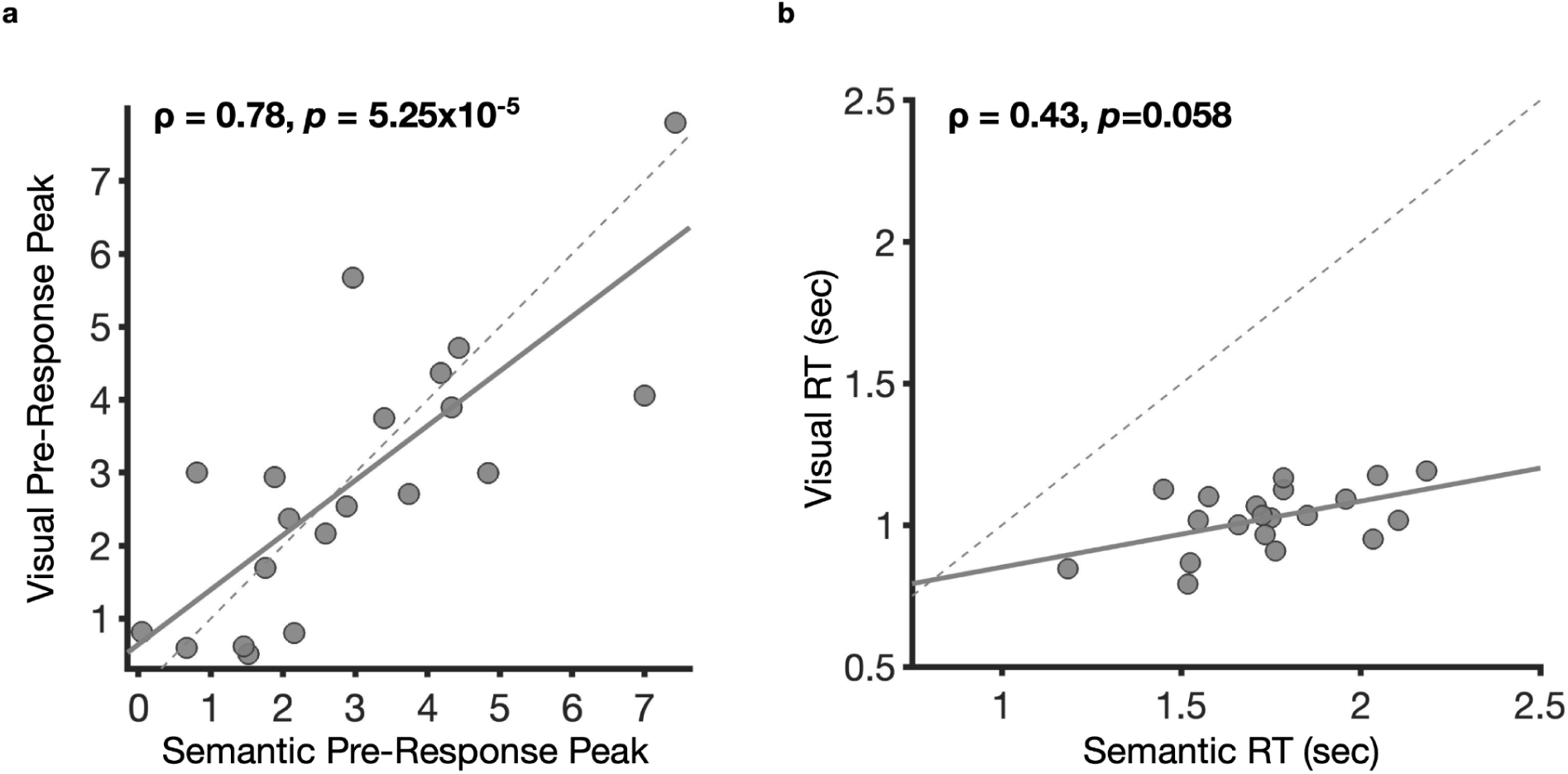
A) Correlation between response-locked semantic memory (x axis) and perceptual (y axis) pre-response peak CPP amplitudes (-100 to-25 ms) collapsed across difficulty levels in Experiment 1 (N = 20). Participants with higher pre-response peaks in the semantic memory task also showed higher pre-response peak amplitudes in the perceptual task, suggesting the possibility of a domain-general evidence accumulation threshold. B) Correlation between response-locked semantic memory (x axis) and perceptual (y axis) median RTs collapsed across difficulty levels, suggesting a common influence on decision speed.

### The CPP is Sensitive to Episodic Memory Strength

We next aimed to extend the findings of Experiment 1 and provide further support for the CPP as a domain-general neural index of evidence accumulation by examining whether CPP dynamics generalized to episodic memory-based decisions. To this end, we conducted a second experiment investigating evidence accumulation during word recognition judgments. If the CPP reflects a domain-general neural signal, its buildup dynamics should mirror those observed in Experiment 1. Evidence strength in Experiment 2 was manipulated by varying the exposure frequency (1x, 2x, or 4x) of words presented during a learning phase. Memory for learned items was subsequently evaluated in a 2AFC test (Figure 6A), and only EEG data from the test was analyzed. As shown in Figure 6B, less exposure during learning led to lower accuracy, *F*(2,50) = 110.2, *p* = 4.72×10^-19^; lower confidence, *F*(2,50) = 51.38, *p* = 7.48×10^-13^; and slower RTs, *F*(2,50) = 67.98, *p* = 8.56×10^-15^ at test (one-way repeated-measures ANOVA).

**Figure 6.**
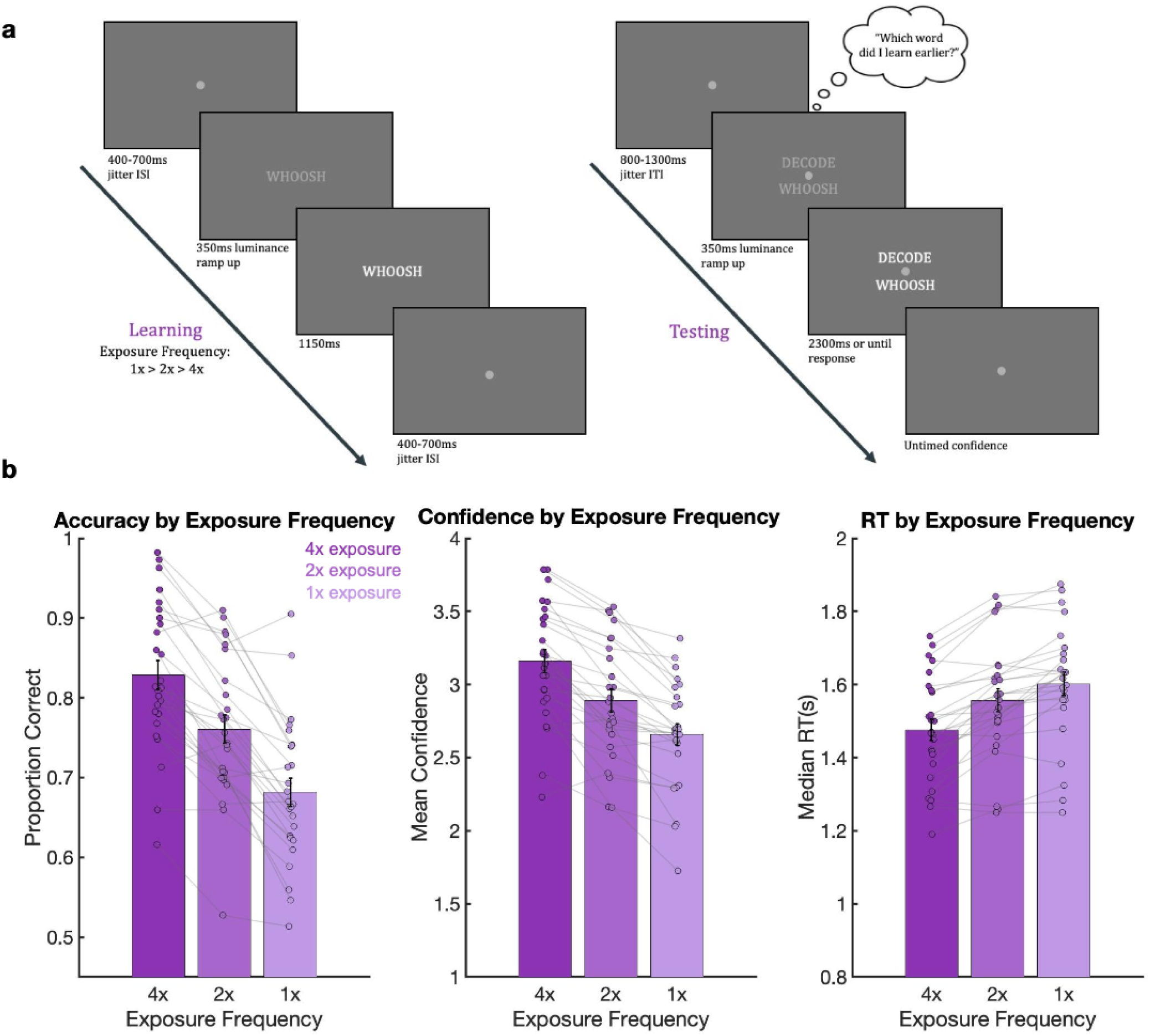
Example trials and behavioral results from Experiment 2. A) Each learning trial (left) began with a central fixation dot and 350ms gradual luminance ramp up of the stimuli, which were then displayed for an additional 1150ms, followed by a randomly jittered inter-stimulus interval between 400-700ms. Stimuli were presented serially at one of three randomized exposure levels (4x, 2x, 1x). Each testing trial (right) began with a central fixation dot and 350ms gradual luminance ramp up of the target–foil word pairs, which were displayed until response (up to a maximum of 2300ms). Participants were tasked with choosing the previously learned word and rating their confidence in that decision (on a scale of 1-4). B) The mean accuracy (left plot) at testing decreased as exposure frequency in learning decreased, mean confidence rating (middle plot) decreased as exposure frequency decreased, and median response time (right plot) increased as exposure frequency decreased. Error bars denote ±1 SEM (across participants). Colored dots over each bar represent individual subjects.

During decision formation, we observed a positive ERP component over central parietal electrodes which peaked around 1,050 ms poststimulus (Figure 7, left) and ramped up prior to motor decision response (Figure 7, right), aligning with our findings in Experiment 1 and resembling the CPP (O’Connell et al., 2012).

**Figure 7.**
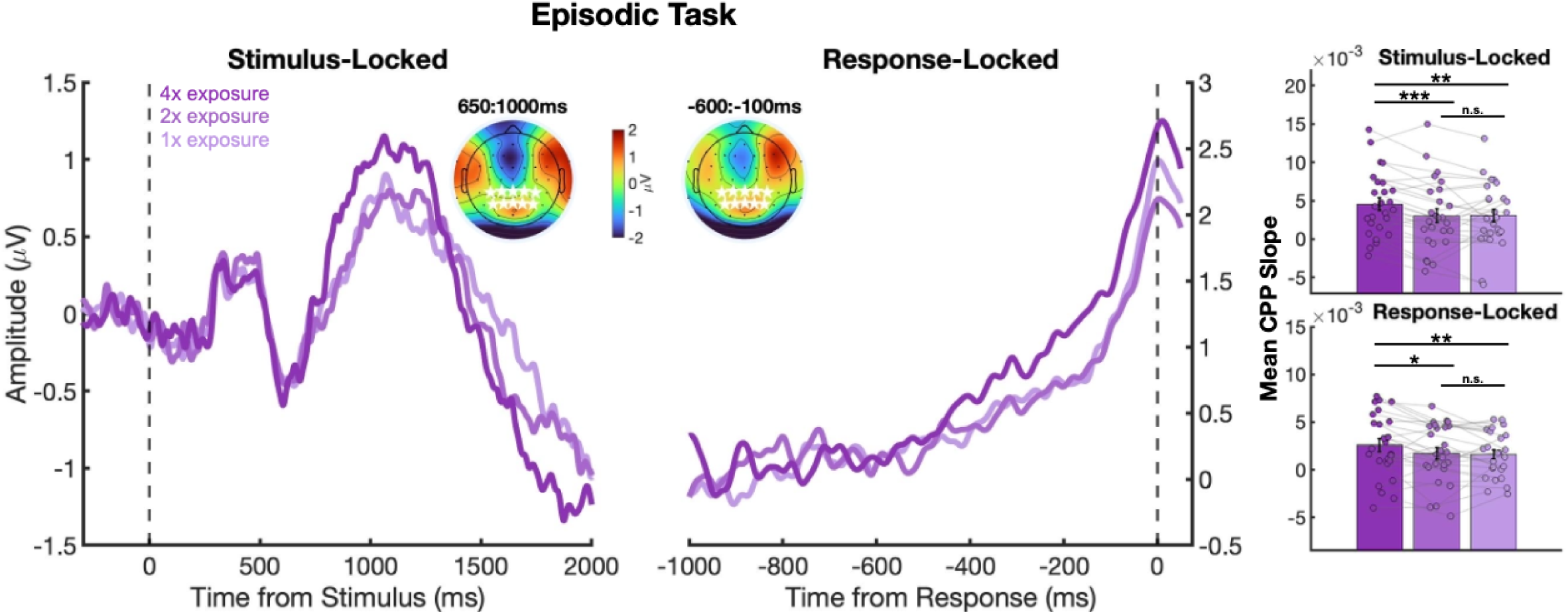
Effects of exposure frequency on the CPP component in Experiment 2. Grand average CPP waveforms were aligned to both stimulus onset (left column) and response execution (right column). The vertical dashed lines denote the time of stimulus onset (left) and time of response execution (right). The inset topographies of the ERP measured after stimulus onset show a positive central-parietal component (electrodes used for analysis are shown in white stars) across both stimulus and response-locked windows. Stimulus-aligned and response-aligned CPP slopes both increased as episodic memory strength increased. Bar plots (far right) show mean CPP slopes per exposure group, with individual subjects plotted as points.

To test if the CPP was sensitive to the strength of episodic memory evidence, we ran two one-way repeated-measures ANOVAs predicting CPP slope from Exposure Frequency (1x, 2x, or 4x) separately for response-and stimulus-locked waveforms. As shown in Figure 7, CPP slopes varied as a function of exposure, steeper buildup rates for more frequently exposed words in both stimulus-locked (*F*(2,50) = 8.61, *p* = 6.11×10^-04^) and response-locked (*F*(2,50) = 4.75, *p* =.013) analyses. Additional *t*-tests were conducted to observe the effects of each exposure level on CPP slope individually, revealing steeper slopes for the 4x condition relative to both 1x (stimulus-locked: *t*(25) = 3.34, *p* =.0026; response-locked: *t*(25) = 2.83, *p* =.009) and 2x (stimulus-locked: *t*(25) = 4.9, *p* = 4.86×10^-05^; response-locked: *t*(25) = 2.38, *p* =.025) conditions. No differences were observed between the 1x and 2x conditions (stimulus-locked: *t*(25) = 0.055, *p* =.96; response-locked: *t*(25) = 0.26, *p* =.79).

### Reaction Time Analyses

To test whether the CPP reflected decision-related behaviors in Experiment 2, we next examined whether CPP slopes varied with RT in the episodic memory task. Trials were categorized as fast or slow in a similar manner to that of Experiment 1. As shown in Figure 8, CPP slopes varied as a function of RT, with faster decisions associated with steeper slopes in both stimulus-locked (*F*(1,25) = 45.43, *p* = 4.60×10^-07^) and response-locked (*F*(1,25) = 11.25, *p* =.0025) analyses. Greater exposure in learning additionally resulted in steeper CPP slopes in both stimulus-locked (*F*(2,50) = 8.72, *p* = 5.61×10^-04^) and response-locked (*F*(2,50) = 4.74, *p* =.013) analyses. A significant interaction between Exposure and RT was observed in both stimulus-locked (*F*(2,50) = 3.80, *p* =.029) and response-locked (*F*(2,50) = 2.79, *p* =.071) analyses, indicating that CPP amplitude varied systematically with both RT and Exposure, and the influence of RT differed across exposure groups. Specifically, the relationship between RT and CPP slope was most pronounced in the lowest exposure condition, where weak prior learning led to shallower accumulation overall and greater separation between fast and slow trials. In contrast, this relationship was attenuated in the highest exposure condition, where stronger prior learning supported more consistent evidence accumulation dynamics and reduced variability in CPP slopes across RT bins.

**Figure 8.**
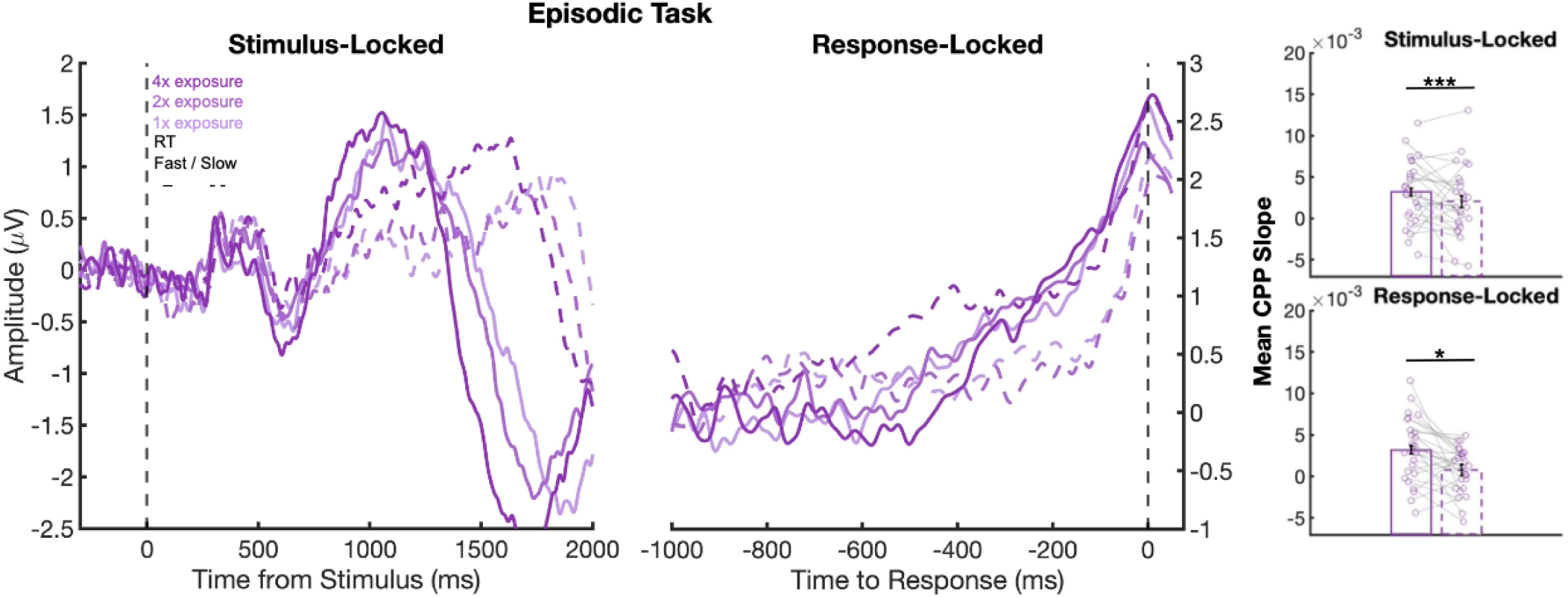
Effects of RT on the CPP in Experiment 2. Grand average CPP waveforms aligned to both stimulus onset (left column) and response execution (right column). The vertical dashed line denotes the time of stimulus onset (left) and time of response execution (right). The CPP was broken down by exposure group and then split into fast (solid line) and slow (dashed line) RT trials within each exposure group based on a median split. Both stimulus-aligned and response-aligned CPP slopes were steeper on fast trials as compared to slow trials.

### Confidence Analyses

As a further test of whether the CPP reflected decision-related behaviors, we next examined whether CPP slopes correlated with subjective confidence. Trials were categorized as low or high confidence in a similar manner to that of Experiment 1. As shown in Figure 9, CPP slopes were positively related with Confidence, such that higher confidence corresponded with steeper slopes in both stimulus-locked (*F*(2,50) = 6.56, *p* =.003) and response-locked (*F*(1,25) = 13.08, *p* =.0013) analyses. Greater exposure during learning resulted in steeper CPP slopes in both stimulus-locked (*F*(1,25) = 8.48, *p* =.0075) and response-locked (*F*(2,50) = 3.99, *p* =.025) analyses. No interactions were observed between Confidence and Exposure in either stimulus-(*F*(2,50) = 0.7, *p* =.50) or response-locked (*F*(2,50) = 0.27, *p* = 0.77) analyses, suggesting that the influence of confidence was consistent across exposure level. This pattern suggests that unlike RT effects, confidence-related modulations of CPP slope were stable across learning conditions, indicating a stable relationship across learning conditions.

**Figure 9.**
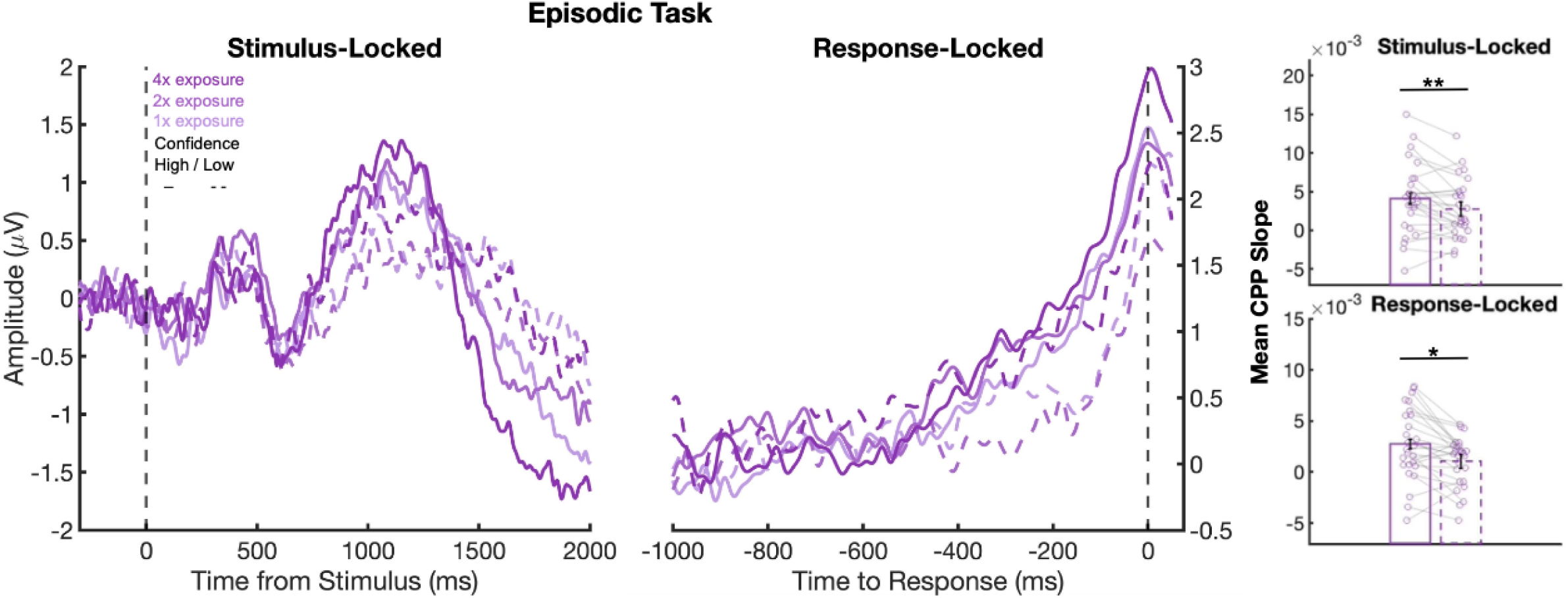
Effects of confidence on the CPP in Experiment 2. Grand Average CPP waveforms aligned to both stimulus onset (left column) and response execution (right column). The CPP was broken down by exposure group and then split into high (solid line) and low (dashed line) confidence trials within each exposure group based on a mean split. Both stimulus-aligned and response-aligned CPP slopes were steeper on high-confidence trials compared to low-confidence trials.

## Discussion

It has been suggested that the decision-making framework of evidence accumulation extends beyond the sampling of externally-driven sensory evidence to include retrieval of internal samples of evidence from memory, though little neural evidence has been presented to support this notion in humans (Ratcliff, 1978; Shadlen & Shohamy, 2016; Van Ede & Nobre, 2024). Across two experiments, we demonstrated that the CPP ERP component associated with tracking evidence accumulation during perceptual decisions similarly tracks evidence accumulation during judgments involving retrieval from semantic (Experiment 1) and episodic memory (Experiment 2). Behaviorally, as expected, easier trials were associated with higher accuracy, faster responses, and greater subjective confidence across all three tasks. At the neural level, we found that the CPP tracks evidence accumulation across perceptual, semantic, and episodic memory decisions with steeper CPP slopes on trials with stronger evidence, faster reaction times, and higher confidence across all tasks.

Our findings contribute to the broader theoretical debate about whether the nature of decision processes are embodied or abstract. Embodied theories propose that cognitive processes are grounded in the body’s sensory and motor systems, such that decision-making may depend on modality-specific representations and actions (Wilson, 2002). In contrast, the observation that the CPP exhibits similar evidence-accumulation dynamics across perceptual and mnemonic domains across two experiments suggests that decision formation can operate abstractly, independent of the evidence source. Prior work has already shown that the CPP can be dissociated from overt motor output (Kelly & O’Connell, 2013; O’Connell et al., 2012), indicating that it indexes an amodal accumulation-to-bound process rather than a motor-specific signal. Extending this logic, our finding that the CPP also tracks evidence derived from internal memory retrieval provides further evidence against a strictly embodied account of decision making. Instead, it supports the view that decision-making relies on shared computational principles across domains, such as evidence accumulation to a common decision threshold, providing a unified decision framework in which a single latent accumulation process underlies diverse forms of choice behavior (Shadlen & Shohamy, 2016).

The fact that similar evidence-sensitive dynamics were observed in our perception and both of our memory tasks suggests that the accumulation of internal evidence over time might be a key computation underlying some judgments based in different cognitive domains. The CPP may therefore reflect the same underlying computation of evidence accumulation, even though the temporal dynamics of accumulation is variable across different sources of decision evidence. In perceptual decisions, evidence accumulation begins shortly after stimulus presentation and continuously builds as long as sensory evidence is available until a decision threshold is reached (Herding et al., 2019; Kelly & O’Connell, 2013; O’Connell et al., 2012; Shadlen & Shohamy, 2016; Twomey et al., 2015). In contrast, because the accumulation of mnemonic evidence depends on cue-driven sampling of stored representations rather than direct access to externally available information, memory decisions could result in greater variability in accumulation onset, slower buildup rates, and longer reaction times than perceptual decisions (Bornstein & Norman, 2017; Ratcliff, 1978; Ratcliff & Starns, 2013; Shadlen & Shohamy, 2016; Weidemann & Kahana, 2016). Such retrieval-dependent variability would likely contribute to the delayed CPP onset and reduced slope observed in the memory tasks, in addition to the overall greater complexity involved in first reading two memory cues before internal evidence sampling can begin.

Notably, with our manipulation of sensory and mnemonic evidence, the effect of difficulty on CPP slopes seemed largely driven by the easy difficulty (Experiment 1) and greatest exposure (Experiment 2) conditions, which clearly differentiated from the medium/2x and hard/1x conditions (but which were not clearly differentiated from one another in either experiment). This pattern of effects mirrors the behavior as the medium/2x and hard/1x conditions have smaller impacts on RT and confidence than the easy/4x condition (see Figures 1B and 6B). This differed from the effects of evidence strength on accuracy, which was more clearly impacted by the strength of evidence at all three levels. Together, this pattern could suggest that the CPP more closely tracks RT and confidence changes across difficulty/exposure levels, rather than accuracy.

One possible concern is that the memory CPP results are in fact driven by perceptual demands related to the reading of word stimuli. While it is true that both memory tasks required participants to read and process word stimuli, the perceptual processing of words themselves was independent of the experimentally manipulated evidence driving the decisions that the CPP slope tracked– namely, the differences in state population (Experiment 1) and exposure frequency (Experiment 2). In other words, while our memory tasks involved perceptual processing, the differences in CPP slopes were specifically modulated by the strength of mnemonic evidence, which was manipulated experimentally and independently of perceptual processing demands.

To the best of our knowledge, the current study is the first of its kind to provide evidence linking the CPP to evidence accumulation in either a semantic or episodic memory task. Recent studies using other mnemonic tasks have identified components reflecting similar decision dynamics in memory. One such component observed in recognition decisions, the Late Positive Component (LPC), has recently been associated with recognition memory evidence strength and ramps up until a recognition decision is made (Sun et al., 2024, 2025; Yang et al., 2019). The LPC seems to scale with RTs on “old” responses as well as distinguish between recollection and familiarity-based responses. Unlike the CPP, however, the LPC is not very clear in stimulus-locked analysis (as opposed to response-locked analysis) and exhibits a left-lateralized topography. Future questions remain about whether the CPP and LPC index the same underlying decision dynamics or reflect different accumulation processes. Of further note, the topography of ERP activity during the CPP time window in Experiment 2 showed additional lateral-frontal components (Figure 7), which were not observed during semantic judgments (c.f. Figure 2). While this could reflect additional brain activity specific to episodic recognition processes, a clear CPP signal was nevertheless observed that was sensitive to evidence strength and predicted decision-related behaviors.

To further test the domain-generality of the CPP, future research could continue to examine the extent to which the CPP may underlie other forms of internal evidence accumulation. For example, whether the CPP encoded evidence accumulation during different types of mnemonic decision tasks, such as free recall or procedural memory, which would extend our findings further across semantic and episodic memory. Additionally, real-world decision-making often requires a combination of novel perceptual stimuli and previously learned stimuli retrieved from memory. As such, it may be of interest to investigate how changes in learned information, expectations, and attentional priorities may shape the temporal dynamics of evidence accumulation. Lastly, it should be noted that our findings so far show a neural signal of evidence accumulation from memory on a trial-and subject-averaged basis. It is possible that on single trials, or for certain subjects, retrieval dynamics do not conform to a gradual build-up of evidence, but rather a more step-like transition in memory access, or myriad other possible temporal profiles of internal evidence access. While our present results do not distinguish between these different accounts of single-trial decision making, this remains an important area for future research, perhaps addressable through joint computational modeling of evidence accumulation dynamics and single-trial EEG activity.

The clear pattern of effects observed across decision modality supports the view that the CPP reflects a shared evidence accumulation mechanism across perceptual, semantic, and episodic decisions. Together, these results indicate that the CPP reflects an abstract neural mechanism underlying decision formation, linking perceptual and mnemonic decisions through shared accumulation dynamics.

## Acknowledgements

We would like to thank Jacob Chaudhry, Jaxon Elsner, Alexandra Mcgowan, Aishwaroopa Narayanan, Grishell Guerrero, and Sonia Murugesh for their help on this project.

